# Differential Wnt/β-Catenin Signaling via TCF7L2/LEF1 Binding Specificity Shapes Cellular and Tumor Phenotypes

**DOI:** 10.1101/2025.11.04.686471

**Authors:** Thomas A. Kluiver, Anna Nordin, Yuyan Lu, Stephanie A. Schubert, Chang Zhang, Xiaochen Duan, Rishi Savur, Marius C. van den Heuvel, Vincent E. de Meijer, Ruben H. de Kleine, Kathelijne Kraal, Ronald R. de Krijger, József Zsiros, Claudio Cantù, Weng Chuan Peng

## Abstract

The mechanisms by which Wnt/β-catenin signaling regulates gene expression in a tissue- and context-specific manner remain poorly understood, limiting our ability to target the aberrant cell growth typical of many Wnt-driven cancers. Here we focus on malignant liver tumors driven by activating *CTNNB1* (β-catenin) mutations that nevertheless display distinct phenotypic states and Wnt outputs. By profiling patient-derived organoids via single-cell transcriptomics and chromatin dynamics, we identify subtype-specific transcriptional and epigenetic profiles. Using CUT&RUN, we show that β-catenin engages distinct genomic regions, dictated by differential association with TCF/LEF family transcription factors. Specifically, we define a novel sequence-specific regulatory element engaged by β-catenin only upon interaction with TCF7L2, revealing that partner choice, independent of *CTNNB1* mutational status, ultimately determines cell fate. Our findings, validated across multiple tumor models and patient tissues, offer a framework for understanding how differential β-catenin-TCF/LEF interaction orchestrates context-specific Wnt signaling outcomes.

**Significance:** Wnt/β-catenin signaling is crucial for development and cancer, yet how it drives different gene programs across tissues is unclear. Using patient-derived liver tumor organoids, we show that β-catenin’s transcriptional output depends on its binding partner: LEF1 or TCF7L2. These factors guide β-catenin to distinct genomic regions, activating either stemness or differentiation genes. We identify a novel helper motif that directs β-catenin-TCF7L2 binding and target selection. By linking partner choice and motif specificity to context-dependent gene regulation, our work provides a unifying mechanism explaining how Wnt/β-catenin signaling produces diverse cellular outcomes.

## Introduction

Wnt/β-catenin signaling is a cell communication mechanism fundamental for regulating cellular proliferation and fate specification during embryogenesis and tissue homeostasis^1,2^. Its mechanism of action centers on WNT ligand-dependent stabilization of cytosolic β-catenin. Once stabilized, β-catenin travels to the nucleus where it forms a complex with a member of the TCF/LEF transcription factor family to regulate Wnt target gene expression. TCF/LEF, by DNA motif recognition, determine the genomic Wnt responsive elements (WREs)^3^. However, how β-catenin regulates different genes depending on the cell type remains an outstanding question in the field.

Dysregulation of Wnt/β-catenin signaling causes developmental abnormalities and many cancers^4,5^. Among cancers, hepatoblastoma offers particularly tractable model to interrogate this pathway in cancer. It is a developmental tumor that is typically driven by activating mutations in *CTNNB1* (encoding β-catenin), with less frequent alterations in other Wnt pathway genes, and otherwise exhibit a very low mutational burden^6,7^. This relatively ‘clean’ genetic background and well-defined phenotypic subtypes provide a powerful framework to dissect how Wnt/β-catenin pathway regulates lineage programs and malignant behavior.

Two main epithelial cell types are commonly observed in hepatoblastoma: the less differentiated ‘embryonal’ histologic subtype (E), characterized by high expression of classical Wnt target genes and associated with worse prognosis^8,9^, and the more differentiated ‘fetal’ histologic subtype (F), which resembles the fetal liver developmental stage^10^. Recent technologies confirmed this classification on a molecular level^6,11–21^. However, it remains unclear how the common genetic driver mutations that stably activate β-catenin leads to different outcomes. This is particularly puzzling given that E and F subtypes often coexist within the same tumor^10^. Thus, liver tumors of developmental origin offer a unique and naturally occurring context to study how overactivation of Wnt/β-catenin signaling could drive distinct cellular outcomes.

Here, we use patient-derived tumor organoids (PDO) representing the two subtypes^20^. By integrating single-cell RNA sequencing (scRNA-seq), single-nucleus ATAC-seq, and CUT&RUN targeting β-catenin and TCF/LEF, we discover that β-catenin physically associates with different genomic elements between E and F cells. We find that the chromatin state and other lineage-specific transcription factors only partially explain this differential β-catenin activity. More surprisingly, we discover that the determinant of β-catenin activity is attributed to the distinct DNA-binding activity of the two main TCF/LEF transcription factors expressed, LEF1 and TCF7L2. LEF1, in the E subtype, is associated with the classic consensus sequence present in well-characterized WREs, while TCF7L2 also recognizes a different DNA motif and drives β-catenin to regulate liver differentiation and metabolism genes. This consensus-mediated genomic association occurs across cell types and tumor models, revealing this as a universal mechanism of Wnt/β-catenin signaling regulation.

Collectively, our study leverages an integrative multi-omics approach in patient-derived organoids to elucidate fundamental aspects of Wnt signaling-driven transcriptional regulation. These insights expand our understanding of Wnt signaling dynamics and establish a robust experimental platform for the investigation of cellular subtype-specific regulatory mechanisms in the context of cancer and developmental biology.

## Results

### Divergent Wnt gene expression programs despite constitutive activation of β-catenin

We employed six unique patient-derived organoid models, representative of E (13E, 17E, 22E) and F subtypes (10F, 13F, 135F) (Fig. 1a)^20^. Among these, 13E and 13F were derived from the same patient (Fig. 1a). Single-cell RNA sequencing (scRNA-seq) performed on these organoids^20^ confirmed a PCA-based separation of the two groups, and presence of distinct cellular features (Fig 1b-d, Supp. Fig. 1a, b). The F subtype expressed high levels of differentiated hepatic markers (*SERPINA1*, *APOA1*, *ALB*) and showed enrichment for gene sets involved in hepatic functions, including xenobiotic metabolism and coagulation (Fig. 1c, d). Conversely, the E subtype expressed high levels of genes associated with stemness (*MSX1*, *PRDM1*, *STMN1*) and epithelial plasticity (*VIM*, *ANXA1*, *MMP16*, *GJA1*) (Fig. 1c). This was accompanied by strong enrichment of the Wnt signaling gene set (Fig. 1d). The differential expression of Wnt/β-catenin related genes was notable (Fig. 1e). The E subtype displayed elevated expression of canonical WNT-associated genes, including *NKD1*, *NOTUM*, *DKK1*, *DACH1*, *MSX1*, *VIM*, and *FGF8*. In contrast, the F subtype showed lower expression of these genes but higher expression of Wnt-regulated metabolic genes (*GLUL*, *OAT*) and additional Wnt targets such as *LGR5*, *ROBO1*, *MYC*, *PROX1* and *SOX9* (Fig. 1e)^22^. These data suggest the presence of a shared Wnt program across both subtypes, as well as subtype-specific enrichment of Wnt target genes.

**Figure 1.**
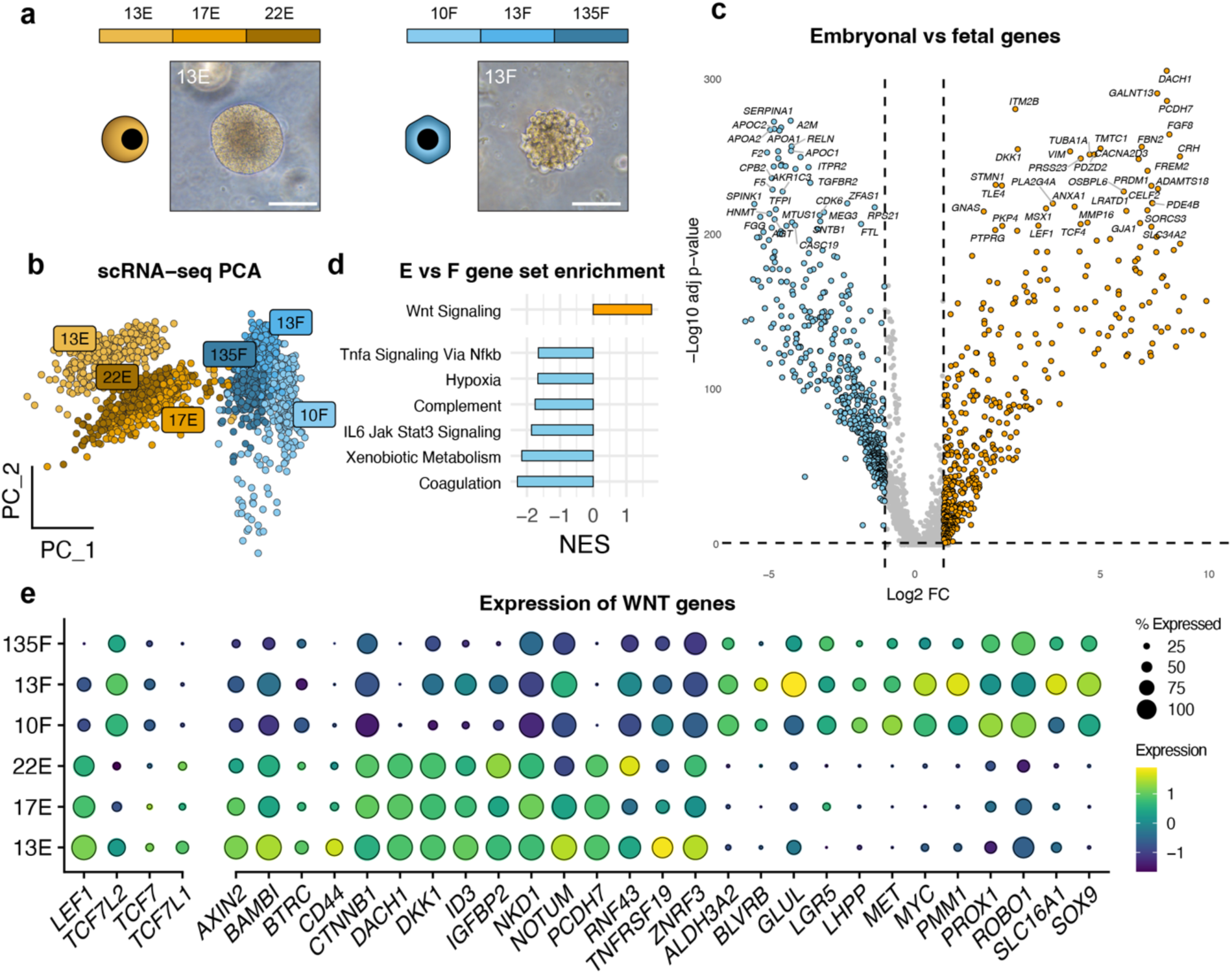
Single-cell transcriptome analysis of PDOs. a. Overview of the six organoid models used in this study, with phase-contrast pictures of representative 13E and 13F organoids. Scale bars, 100 µm. b. scRNA-seq PCA embedding splits embryonal and fetal tumor cells. c. Volcano plot showing differentially expressed genes between the two subtypes. d. Gene set enrichment plot showing enrichment for Hallmark gene sets in both subtypes. NES: normalized enrichment score. e. Dot plot showing expression of Wnt targets and related genes in embryonal (E) and fetal (F) models.

Next, we examined the expression of TCF/LEF transcription factors, the main DNA binding partners of nuclear β-catenin. Among these, TCF7L2, a Wnt effector broadly expressed in the liver and known to regulate hepatic metabolic program^22^, was detected in both E and F subtypes, with higher expression in the F subtype (Fig. 1e, Supp. Fig 1c). In contrast, LEF1, typically associated with a progenitor phenotype and stemness in development and cancer^23^, is specifically upregulated in the E organoids (Fig. 1e, Supp. Fig 1c). Notably, LEF1 is generally absent in mature hepatocytes, and its expression is consistent with a less differentiated, more progenitor-like identity in E organoids. The expression of TCF7 and TCF7L1 was low in both subtypes, indicating a limited role for these transcription factors in hepatoblastoma (Fig. 1e).

### Subtype-specific chromatin accessibility and motif enrichment

To investigate whether the differential Wnt/β-catenin responses observed in E versus F organoids are caused by subtype-specific epigenetic features, we utilized our single-nucleus chromatin accessibility profiling parallel to transcriptomics (scATAC/RNA-seq) dataset featuring four organoid models (13E, 17E, 10F, 13F). Dimensionality reduction using ATAC-based latent semantic indexing (LSI) separated the organoids by subtype, consistent with PCA and UMAP analyses (Fig. 2a, Supp. Fig. 2a), indicating that the two subtypes have different chromatin states. Motif enrichment analysis performed across the sequence underlying the ATAC-seq peaks revealed subtype-specific signatures: E organoids were enriched for motifs of TCF/LEF, p53, and DLX/LHX transcription factors, while F organoids showed motifs of hepatic regulators, such as GATA, FOXA, ONECUT, HNF1A/B, and HNF4A/G (Fig. 2b, Supp. Fig. 2b). Thus, the heterogeneity in chromatin accessibility mirrors that observed via scRNA-seq, indicating distinct transcriptional networks.

**Figure 2.**
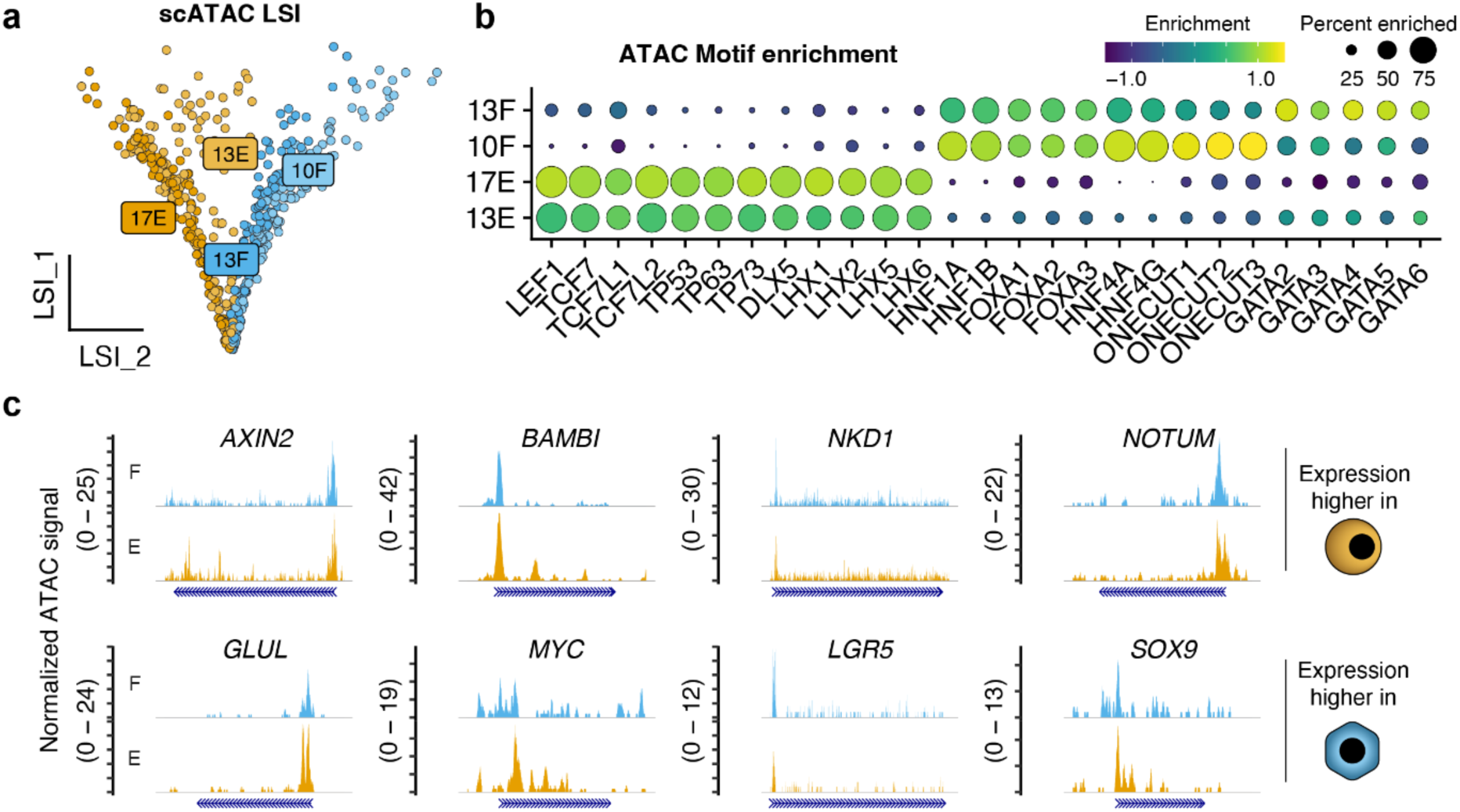
Chromatin accessibility analysis of PDOs. a. snRNA/ATAC-seq LSI embedding of embryonal and fetal tumor cells. b. Dot plot showing ATAC motif enrichment of selected TF families. c. Coverage plots showing ATAC signal at promoter regions of Wnt targets that were overexpressed in either E or F organoids.

We then examined the chromatin accessibility at promoters of Wnt targets enriched in each subtype, including *AXIN2*, *BAMBI*, *NKD1* and *NOTUM* for E organoids and *GLUL*, *MYC*, *LGR5* and *SOX9* for F organoids (Fig. 1e, Fig. 2c). Surprisingly, promoter accessibility was comparable between subtypes (Fig. 2c). Additionally, the F subtype showed increased accessibility at promoters of hepatic genes such as *ASGR1*, *APOA2*, *APOB* and *TF* (Supp. Fig. 2c). Overall, while broader chromatin landscapes and motif enrichment differs between subtypes, promoter accessibility at common Wnt targets appeared comparable.

### β-catenin displays subtype-specific genome-wide binding

We next assessed whether differential recruitment of β-catenin and its TCF/LEF partners to DNA might explain the divergent Wnt signaling outcomes. We mapped genome-wide binding of β-catenin and TCF/LEF family members with CUT&RUN-LoV-U^24^ on the E and F organoids (Fig. 3a, PCA in Supp. Fig 3a). LEF1 and TCF7L2 were the only TCF/LEF factors yielding reliable signal due to rheir higher expression pattern. After quality control we merged replicates, called peaks using GoPeaks (adj. p<0.05) against the matching IgG negative controls, and compiled a set of “Wnt peaks” for each subtype: loci bound by β-catenin together with LEF1 or TCF7L2 in at least two of the three organoid lines (Fig. 3a, Supp. Fig. 3b, Supp. Table 1). In addition, assay validity was confirmed by robust β-catenin binding at WREs within the *AXIN2* locus in both subtypes (Fig. 3b, center).

**Figure 3.**
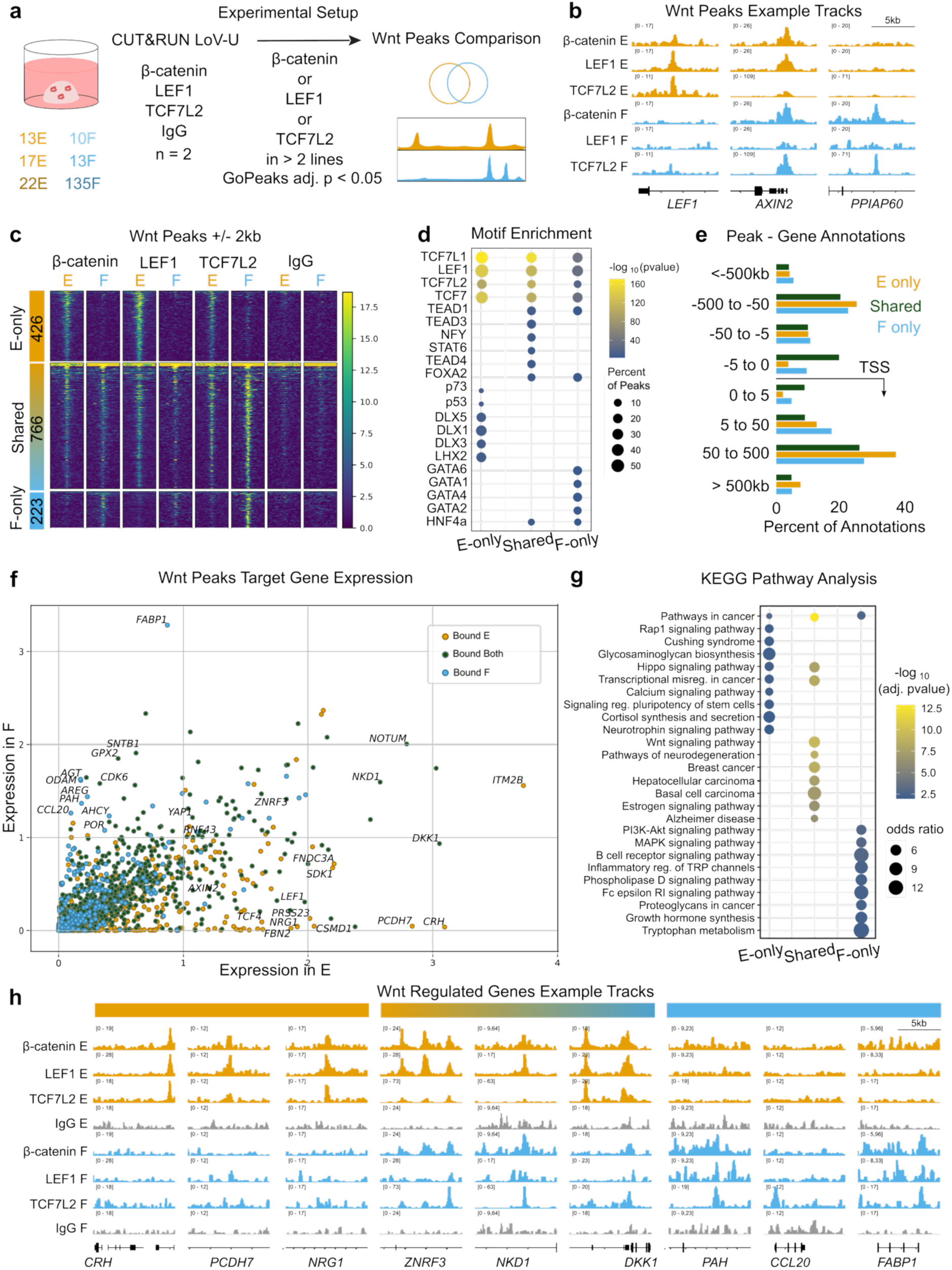
CUT&RUN analysis reveals subtype-specific Wnt/β-catenin binding activity. a. Experimental design overview and the definition of Wnt peak sets. b. Example genome browser visualization of Wnt peaks specific to embryonal lines (*LEF1,* left), common to both (*AXIN2*, center) and specific to fetal lines (*PPIAP60*, right). For genome browser visualization, normalized bigwigs from all lines from each subtype and target were averaged. c. Heatmaps of signal centered within peak regions and showing +/- 2kb for all CUT&RUN targets over E-only peaks (top, 426), shared (middle, 766) and F-only peaks (bottom, 223). Heatmaps are based on the averaged bigwigs. d. Motif enrichment dot plot of HOMER identified known motifs in E-only, shared and F-only Wnt peak regions, showing high enrichment of TCF/LEF consensus sequences as well as subtype-specific motifs. Top 10 motifs are shown, cutoff q < 0.05, dot color represents significance and dot size percentage of peaks. e. Peak-gene annotation frequencies according to GREAT analysis, showing a higher instance of promoter occupancy in shared Wnt peaks. f. Graph showing expression in E (x-axis) versus expression in F (y-axis) for all annotated Wnt target genes. Dots are colored based on bound status. g. KEGG pathway enrichment dot plot based on expressed target genes bound only in E, bound in both, or only in F. Top 10 results are shown, cutoff adj. p < 0.05. Dot color represents significance and size represents odds ratio. KEGG analysis was done with all expressed genes as a background set. h. Additional genomic loci of E-only targets (left), shared Wnt target genes (center) and F-only targets (right).

About half of the peaks (766) were shared between E and F, while 426 were E-only and 223 F-only, revealing a conspicuous divergence in the physical activity of nuclear β-catenin (Fig. 3b, c, Supp. Table 1). Motif enrichment analysis revealed that all three groups of peaks (shared, E-only, F-only) were enriched for TCF/LEF consensus (Fig. 3d). In addition, in the E-only set we detected motifs of p53 and members of the DLX homeobox family (DLX1/3/5), while in F-only, consistent with a more differentiated phenotype, we found motifs of GATA and HNF4A, transcription factors associated with hepatocyte differentiation and metabolic regulation. These motifs match those identified in the scATAC-seq and indicate that β-catenin targeted genomic regions are not universally predetermined but change even across closely related cellular subtypes.

E and F shared peaks were predominantly located close to promoter regions (28% within 5Kb of the TSS), while subtype-specific peak regions are more gene-distant (only 5% of E-only and 13% of F-only are within 5Kb of the TSS; Fig. 3e). This indicated that one primary differentiator in β-catenin’s activity is the engagement with subtype-specific enhancers. Intersecting CUT&RUN peaks with subtype-specific differential gene expression allowed us to define the biological processes directly driven by β-catenin (Fig. 3f, g, Supp. Table 2). Shared targets like *ZNRF3*, *RNF43*, *NKD1*, and *DKK1* confirmed universal activation of Wnt signaling (Fig. 3f, h). E-only targets included genes for aggressive growth and invasiveness (*PCDH7, NRG1, FNDC3A, SDK1*), implicating β-catenin in the regulation of the enhanced metastatic potential and therapy resistance of the E subtype (Fig. 3f, h). F-only targets included metabolic genes such as *PAH*, *FABP1* and *POR,* emphasizing that β-catenin fosters hepatic differentiation and metabolic specialization of the F subtype (Fig. 3f, h). KEGG pathway enrichment analysis, using all expressed genes as background, confirmed this pattern: shared β-catenin targets were associated with cancer pathways and Wnt signaling, E-only targets were enriched for stem cell pluripotency genes, and F-only targets for metabolic pathways and growth hormone synthesis (Fig. 3g). These data demonstrate that β-catenin contributes to orchestrate differential gene expression programs even in related cellular subtypes within the same tumor.

### Chromatin accessibility alone does not explain differential β-catenin binding patterns

A plausible explanation for the presence of E- and F-only β-catenin peaks could be differential chromatin accessibility in the two subtypes. We explored the scATAC-seq data focusing on Wnt peaks as defined in Fig. 3. In F organoids, the genomic regions corresponding to E-only peaks were mostly, yet with some exceptions, characterized by low scATAC-seq signal, suggesting that the vast majority of the underlying regulatory elements are not available for regulation by β-catenin in this context (Fig. 4a). However, the E organoids displayed comparable open chromatin in the genomic regions corresponding to F-only peaks relative to F organoids (Fig. 4a). Despite being accessible, the F-only regions are not actively engaged by β-catenin or TCF/LEF in E organoids. This indicated that chromatin accessibility, particularly in E organoids, and in a subset of loci in F organoids, do not fully account for the differential binding of β-catenin.

**Figure 4.**
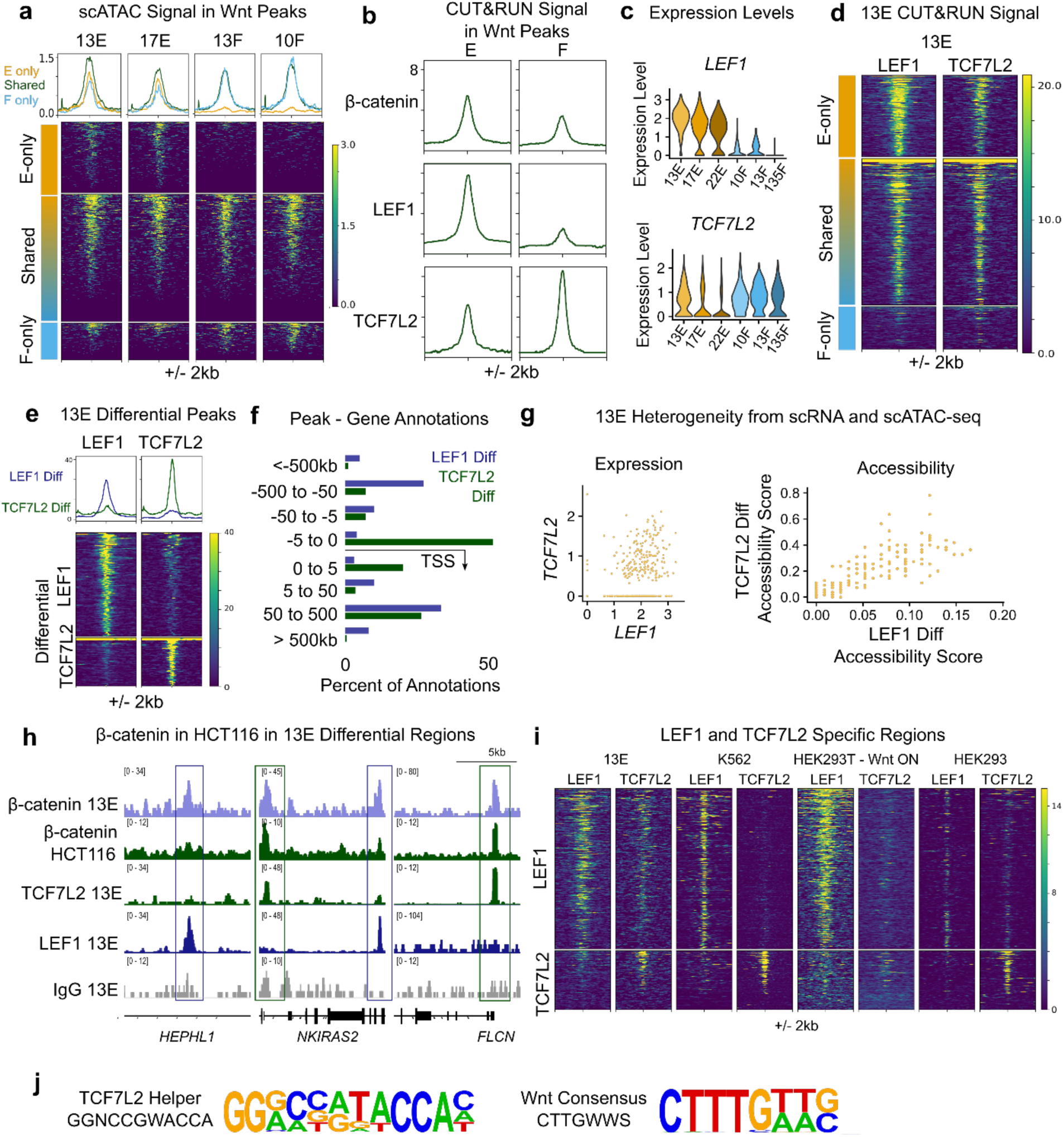
Differential activity of LEF1 and TCF7L2. a. Chromatin accessibility scATAC-seq signal from 13E, 17E, 13F, and 10F plotted within Wnt peak subsets. E-only regions become inaccessible in F lines, while shared and F-only regions remain comparably accessible in both subtypes. b. Signal profiles of β-catenin, LEF1, and TCF7L2 within Wnt peaks, showing the median profile. In E-lines (left, average of all three lines shown), β-catenin and LEF1 signal are higher than TCF7L2 signals, versus in F-lines (right), TCF7L2 signal is strongest. c. *LEF1* and *TCF7L2* expression levels in the organoid lines according to scRNA-seq. Most E-lines expression predominantly *LEF1*, and F-lines *TCF7L2*, except for 13E which expresses both. d. Signal profiles of LEF1 and TCF7L2 CUT&RUN in 13E within Wnt peak regions. e. Differentially occupied regions between LEF1 and TCF7L2 according to PePr (p<0.00005, FC>2) in 13E. f. Peak-gene annotation via GREAT revealed that LEF1 differential regions are located distal to genes, versus TCF7L2 differential regions are predominantly close to the TSS. g. Left: scatter plot showing *LEF1* versus *TCF7L2* expression, revealing that most cells express either only *LEF1*, or both, and there are not subpopulations that express only *TCF7L2.* Right: scatter plot of average accessibility score within LEF1 differential regions versus TCF7L2 differential regions. The correlation indicates these regions are typically equally accessible within single cells, and do not represent subpopulations with the 13E line. h. Genomic browser example views of LEF1 and TCF7L2 differential regions for β-catenin CUT&RUN in HCT116 and data from 13E. i. Heatmaps of LEF1 and TCF7L2 specific regions identified in at least 2 of 4 matched datasets for each protein. j. Motif logos for the identified TCF7L2 helper site and the Wnt consensus motif.

### Locus-specific partnership between β-catenin and TCF/LEF transcription factors

We noticed that the CUT&RUN average signal of the two expressed TCF/LEF factors in PDOs, LEF1 and TCF7L2, depended on cellular subtype: in E cells, TCF7L2 signal was weaker than both LEF1 and β-catenin signals, while in the F lines this ratio is reversed and TCF7L2 was consistently the strongest (Fig. 3c, 4b). This pattern follows the switch in expression: E cells express higher *LEF1* which decreases in F cells in favor of *TCF7L2*, (Fig. 4c), akin to hepatocyte differentiation in the liver^25^. The LEF1-to-TCF7L2 expression switch is accompanied by a switch in genomic loci recognized and consequently bound by β-catenin (Fig. 3c). Therefore, it might be the different activities of LEF1 and TCF7L2 that confer locus-specific association to β-catenin. However, we could not exclude that additional context-specific cofactors drive the different genomic association of LEF1 and TCF7L2 in E versus F, and these Wnt effectors would be redundant were they expressed in the same cells.

The decisive test for this came from the 13E organoid line, which robustly expresses both *LEF1* and *TCF7L2*. Accordingly, this line displays comparable LEF1 and TCF7L2 binding signal (Fig. 4d). We used PePr^26^ to determine regions differentially bound by either LEF1 or TCF7L2 within the 13E line (p<5 x 10^-5^, FC>2, Supp. Table 3) and identified 131 regions where β-catenin partners only with LEF1, and 61 regions where it partners only with TCF7L2 (Fig. 4e). LEF1-only regions were predominantly gene-distal, likely functioning as enhancers, whereas TCF7L2-only peaks were mainly near the TSS, suggesting that β-catenin/LEF1 and β-catenin/TCF7L2 complexes preferentially act on enhancers and promoters, respectively (Fig. 4f). Of note, both the LEF1-only and TCF7L2-only peaks showed comparable enrichment of β-catenin CUT&RUN signal (Supp. Fig. 4a), lending credibility to these peak sets, and indicating that both must be considered as genuine Wnt/β-catenin target genes. Finally, scRNA-seq data of 13E cells shows that while some cells express only *LEF1*, most co-express *LEF1* and *TCF7L2* (Fig. 4g, left). Importantly, scATAC-seq data showed that there are no cells which have open chromatin in exclusively LEF1-only or TCF7L2-only peaks (Fig. 4g, right). These data collectively show that each β-catenin molecule is directed to specific loci predetermined by which TCF/LEF it is associated to.

### Universal association of LEF1 and TCF7L2 on different Wnt-responsive elements

We explored multiple datasets from us and others to test whether the skewed binding pattern of β-catenin towards the same LEF1-only and TCF7L2-only differential regions was universal. The HCT116 colorectal cancer line, which also carries activating mutations in *CTNNB1*, is a good test case of primary regulation by TCF7L2^27^. In these cells, β-catenin consistently binds to the majority of TCF7L2-only regions (45 of 61, 74%), but only to a minority of LEF1-only peaks (29 of 131, 22%). This pattern was confirmed by signal heatmaps and inspection of several loci, showcasing a β-catenin profile in HCT116 that largely matches that of the β-catenin/TCF7L2 complex in 13E organoid (Fig. 4h, Supp. Fig. 4b). It is important to point out that HCT116 cells do not express LEF1, and TCF7L2 appears incapable of fully recapitulating LEF1 binding profile, confirming the selective nature of their different modes of action and the unexpected, limited redundancy. We also found several matched datasets of other cell types expressing both LEF1 and TCF7L2, including our previous CUT&RUN data in CHIR99021-stimulated HEK293T cells (Wnt-ON)^24^, and ChIP-seq datasets in HEK293 and K562 cells from ENCODE^28,29^. Inspection of β-catenin signal across the Wnt peaks seems to respect the pattern identified in hepatoblastoma across all the models considered. (Supp. Fig. 4c).

To identify cell-type agnostic sets of LEF1-only and TCF7L2-only peaks, we overlapped the peaks between the four matched datasets (13E, HEK293T Wnt-ON, HEK293, K562) for LEF1 and TCF7L2, and considered those peaks present in at least 2 different models. This yielded 2220 LEF1 and 983 TCF7L2 peaks. Of these, only 328 (ca. 10% of total) were shared between the two factors. We pinpointed 964 LEF1-only peaks (never called as TCF7L2 peaks; 0 of 4 datasets), and 354 TCF7L2-only peaks (never called as LEF1 peaks; 0 of 4 datasets). Contrary to the expected large redundancy in DNA-binding capabilities, our data identified considerable regions that are only bound by one of the two Wnt regulators (Fig. 4i, Supp. Table 3).

### A novel TCF7L2 long-helper motif directs genome-wide binding specificity

Our analyses showed that that neither expression pattern, nor chromatin accessibility could explain the differential binding of LEF1 and TCF7L2. We tested whether this could be due to an intrinsic difference in their DNA-binding capability. *De novo* motif analysis on the LEF1-only and TCF7L2-only peaks revealed that while LEF1-only peaks were enriched for the TCF/LEF consensus (match score 0.95), the TCF7L2-only binding sites were characterized by an uncharacterized 5’-GGNCCGWACCA-3’ consensus sequence with no convincing match (top ranked was that of the PRDM family with a low score of 0.68) (Fig. 4j). This motif was almost identical to the top motif identified in the 13E TCF7L2 differential peaks (Supp. Fig 4d), and it contains, but it is not limited to, the RGGC helper motif bound by the C-clamp domain, present in TCF7 and TCF7L2^30–33^. This motif is longer than and does not fully match the 5’-GCCGCCR-3’ helper site for TCF7 previously identified via genome-wide analysis in DLD1 cells by the Waterman lab^34^.

We searched across the LEF1-only and TCF7L2-only peaks for the classical TCF/LEF motif: 5’-CTTTGWWS-3’, the short-helper: 5’-RCCG-3’, and the new TCF7L2 long-helper: 5’-GGRCCRTACCAC-3’. We found that the LEF1-only peaks are enriched for multiple TCF/LEF motifs with the same peak (obs/exp 1.55, logP -18.3), but not for co-occurrence of the TCF/LEF motif with either of the two helper sequences (obs/exp 0.94 and 0.74 for short helper, and TCF7L2 helper, respectively) and can be thought of as classical WREs. The TCF7L2-only peaks, on the other hand, were only mildly associated with repeats of the classic TCF/LEF motif (obs/exp 1.12, logP - 2.58) and were also not characterized by co-occurrences of the classic motif with either helper sequence (obs/exp 1.0, 0.57, respectively). Instead, they were enriched for co-occurrence of multiple TCF7L2 long-helper sequences (obs/exp 1.68, logP -2.68), but not for co-occurrence of the short-helper (obs/exp 0.96), despite this is contained within the TCF7L2 long-helper. This indicated that the TCF7L2 long-helper motif is a functional unit that allows differentiation of binding sites specific for TCF7L2 from LEF1. Whether the long-helper might differentiate TCF7L2 from TCF7 that primarily uses the short-helper remains to be determined. These results unearth a new long-helper recognition sequence for TCF7L2, which, even in the absence of a classical TCF/LEF motif, may facilitate DNA binding and Wnt target gene regulation.

### TCF7L2 helper motif mediates a hepatic-lineage gene program

To investigate the gene expression programs potentially regulated by TCF7L2, we performed combined scRNA/ATAC-seq on tumor tissues of patients 13 and 17 (from which organoids 13E/F and 17E were derived, respectively). After filtering low quality cells and selecting for tumor cells, we identified clusters corresponding to E and F subtypes in both samples. Data were integrated using Harmony^35^ and PCA revealed a separation of the E and F subtypes along the first principal component (Fig. 5a). The two subtypes in patient tumors expressed many of the same markers observed in our organoid dataset (Supp. Fig. 5a, b). Notably, this included a clear enrichment of expression of *LEF1* in the E subtype, *HNF4A* in the F subtype, and *TCF7L2* in both (Fig. 5b).

**Figure 5.**
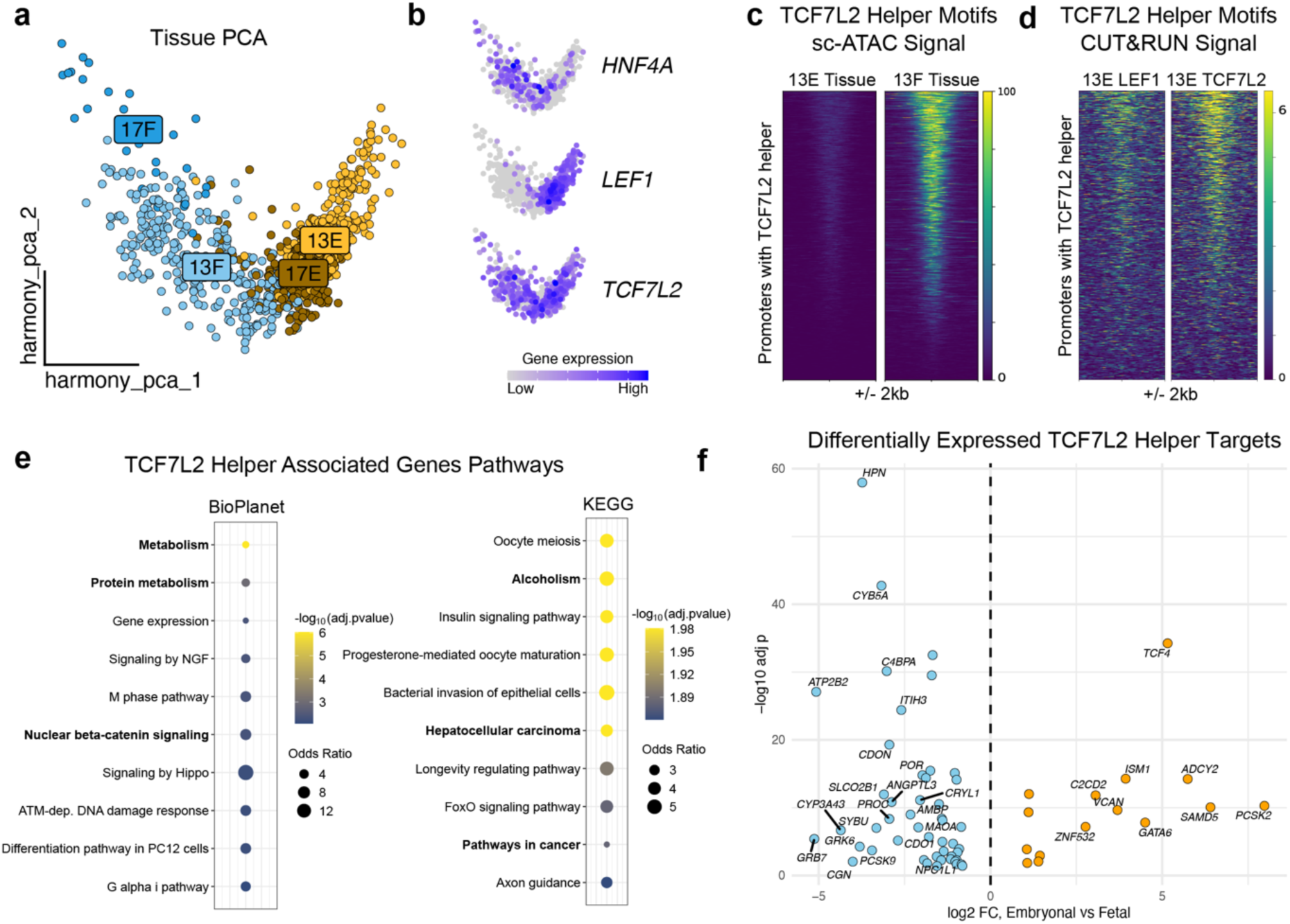
TCF7L2 directs a hepatic lineage gene program. a. Integrated PCA plot of tumor cells from patient tissues. b. Gene expression of *HNF4A*, *LEF1* and *TCF7L2* plotted on top of the integrated PCA plot. c. scATAC signal of 13E and 13F tissue cells within promoter regions containing TCF7L2 helper motifs, showing higher accessibility in 13F. d. CUT&RUN signal for LEF1 and TCF7L2 from 13E organoids over promoter regions containing TCF7L2 helper motifs, showing higher signal in TCF7L2 compared with LEF1. e. Pathway enrichment analysis for genes associated with TCF7L2 helper motifs. Both BioPlanet (left) and KEGG (right) databases result in enriched terms associated with the more differentiated hepatic phenotype. f. Volcano plot showing differentially expressed TCF7L2 helper motif-associated genes within the tissue cells.

Next, we searched the open chromatin regions within the tissue for occurrences of the TCF7L2 long-helper motif. Of 2,752 motif instances, 844 (31%) were located within 5kb of a TSS (Supp. Table 4). These promoters were more accessible in F versus E cells within the tumors (Fig. 5c). We plotted the CUT&RUN signal from the organoids within these TSS-adjacent regions and confirmed that they contain strong TCF7L2 signal enrichment and comparably weak LEF1 signal, validating the usage of this motif to identify differential targets of these two factors (Fig. 5d). When we explored the genes associated with these motifs, the highest enriched KEGG and BioPlanet pathways included hits like hepatocellular carcinoma (adj. p 0.01, odds ratio 3.67) and metabolism (adj. p 9.7 x 10^-7^, odds ratio 2.35), terms associated with a more differentiated F phenotype, and interestingly even nuclear β-catenin signaling (adj. p 0.005, odds ratio 6.32) (Fig. 5e). TCF7L2 helper motif-associated genes differentially expressed in F versus E cells included key hepatic targets, such as *HPN*, *CYP3A43*, *PCSK9* and *POR* (Fig. 5f). Our data suggest that DNA sequence recognition by different TCF/LEF family members is a primary mechanism for the divergent Wnt/β-catenin nuclear activity across different cell types, driving phenotypic differences.

## Discussion

One fundamental unresolved question in developmental biology is how signaling pathways, typically represented as linear series of events culminating in a pathway-specific transcriptional complex, can regulate gene expression in a tissue- and context-specific manner. The Wnt/β-catenin pathway exemplifies this complexity, and our inability to define its cell-type specific mechanisms hinders the identification of inhibitors that could target specific Wnt-driven cancer, without perturbing homeostatic Wnt signaling.

Here we leverage hepatoblastoma, a developmental liver cancer, as a unique model to study Wnt/β-catenin regulation given the presence of two distinct embryonal (E) and fetal (F) phenotypes within a tumor, despite sharing the common causative activating mutations in *CTNNB1*. The patient-derived organoid lines we employ preserve the E and F phenotypes and their defining transcriptional features. Single-cell transcriptomics combined with chromatin accessibility profiling provided reference datasets that reveal regulated gene expression modules, in which lineage-specific transcription factors drive the two distinct cell fates. To uncover the direct causative role of β-catenin into these genetic programs, we mapped binding sites of β-catenin and the related TCF/LEF factors with CUT&RUN, across all the models. This revealed that β-catenin exhibits subtype-specific activities aligned with their divergent fates, which is not explained by *CTNNB1* mutational status alone.

Our key finding, that the E and F subtypes are characterized, and perhaps defined, by a switch in *LEF1* to *TCF7L2* expression, has important implications. These two factors, while being largely redundant, have also been hinted to possess different functions. LEF1 acts as a transcriptional activator within progenitor-like contexts and was often associated with promoting epithelial-to-mesenchymal transition (EMT), stemness, and invasiveness in cancer^23^. In contrast, TCF7L2 was shown to mediate differentiation, notably maintaining proliferation and metabolic function in tissues such as the intestinal epithelium and liver^22^.

Our findings reveal a clear distinction between these two transcription factors, showing that they direct β-catenin to regulatory elements in a locus-specific manner. This difference is caused by recognition of a previously uncharacterized motif on the DNA, bound by TCF7L2, and different from the classic WREs with which LEF1 mostly associates. TCF7L2 is one of the two TCF/LEF, in vertebrates, to possess the additional C-clamp domain that anchors it to WRE via an additional helper motif. Our identification not only sets TCF7L2 apart from the previous motif defined for TCF7^34^ but points to a TCF7L2 regulation of target genes that seems largely independent by the presence of classic WREs. This is surprising, as it expands the definition of Wnt target genes and shows that TCF/LEF might be more different than previously thought in their DNA binding capability. We find surprising that it is the regulatory syntax of DNA elements, in addition the accessibility of the sites, and the association with tissue-specific regulators, that defines subtype-specific Wnt/β-catenin targets. This is determined by the choice of TCF/LEF partner, a family of proteins historically viewed as largely redundant and distinguished mainly by tissue-specific expression patterns. Here, we found that even when both LEF1 and TCF7L2 are simultaneously present in individual cells, they can recruit β-catenin to their own distinct regulatory elements. In addition, we showed that this differential motif usage contributes to the establishment of distinct cellular and tumor phenotypes, even in related cell-types that co-exist within the same tumor.

An important aspect that remains to be determined is the cooperation between the LEF1/β-catenin and TCF7L2/β-catenin complexes with lineage-specific transcriptional regulators. Our analysis suggests that such interaction might be an important contributor to their activity. The LEF1/β-catenin and TCF7L2/β-catenin complexes, in fact, drive expression of lineage-specific genes by binding to regulatory regions that carry motifs for lineage-determining transcription factors. A prominent example is HNF4A, a master regulator of hepatic differentiation, whose motifs are particularly enriched in the F subtype, which is characterized by a more differentiated phenotype closer to adult hepatocytes. While several studies identified an interplay between β-catenin and HNF4A, a closer inspection into their shared genomic action and dependency is much needed to reveal new regulatory logic underlying Wnt-driven differentiation programs.

## Data availability

Raw sequencing data have been deposited in the European Genome-phenome Archive (EGA) and will be made public upon publication.

## Author Contributions

T.A.K. and A.N.: Methodology; Formal analysis; Investigation; Data curation; Visualization; Writing (equal contribution), Y.L., C.Z., S.A.S., X.D., and R.S.: Investigation. M.C.v.d.H., V.E.d.M., R.H.d.K., K.K., and J.Z.: Resources. R.R.d.K.: Investigation; Resources. C.C. and W.C.P.: Conceptualization; Methodology; Supervision; Funding acquisition; Writing (co-senior authors).

## Acknowledgements

We would like to extend our gratitude to the Máxima Comprehensive Childhood Cancer (M4C) Liver Tumor group, which includes Martine van Grotel, Kees van de Ven, Lideke van der Steeg, Anneloes Bohte, Annemieke Littooij, Maarten Smits, Marijn Scheijde-Vermeulen, Liset Lansaat and Charlotte van Aart. We would also like to extend our gratitude to the Princess Máxima Center Single-Cell Genomics Facility (Thanasis Margaritis, Philip Lijnzaad, Tito Candelli, Aleksandra Balwierz), Big Data Core and the Kemmeren group. Research reported in this publication was supported by Oncode Accelerator, a Dutch National Growth Fund project under grant number NGFOP2201, and by Kinderen Kankervrij (KiKa) Project 425. We are grateful for the generous donations from the public through the Princess Máxima Center Foundation (Kus van Kiki). The Princess Máxima Center Single-Cell Genomics Facility is supported by KiKa. Work in the Cantù lab is supported by Cancerfonden (21 1572 Pj and 24 3487 Pj), the Swedish Research Council, Vetenskapsrådet (2021–03075 and 2023-01898), Additional Ventures (USA) (SVRF2021-1048003), Linköping University support and LiU/RÖ-Cancer. Sequencing of CUT&RUN samples was performed at the Core Facility at the Faculty of Medicine and Health Sciences, Linköping University. C.C. is a Wallenberg Molecular Medicine (WCMM) and SciLifeLab Fellow and receives generous financial support from the Knut and Alice Wallenberg Foundation.

## Methods

### Organoid culture

Six organoid lines (embryonal 13E, 17E, 22E and fetal 10F, 13F and 135F), were profiled and cultured as in Kluiver *et al.*^20^ Organoids were maintained in liver tumor medium (Advanced DMEM/F-12 plus B27, N2, N-acetyl-l-cysteine, gastrin, EGF, 10% RSPO1 CM, HGF, nicotinamide, A83-01, forskolin, Y-27632 and next-generation WNT surrogate) as described previously.

### Organoid single cell data analysis

Organoid scRNA-seq and snRNA/ATAC-seq datasets from Kluiver and Lu *et al.* were reanalyzed in R 4.4.0, and only data from the six models used in this study were retained. Analysis was performed using Seurat^36–38^ (v5.1.0). For the scRNA-seq dataset, cells with 2-20% mitochondrial RNA were retained; the top and bottom 10% of nFeature_RNA per line were excluded, and all six lines were downsampled to 300 cells per line. Data was processed using SCTransform; where mitochondrial, cell-cycle and cell-cycle correlating genes were removed from the variable features set as in Kluiver *et al*.^20^ Dimensionality reduction used PCA and UMAP (5 dims). Differential expression between embryonal and fetal groups applied FindMarkers (min.pct = 0.5). The RNA/ATAC multiome analysis employed Seurat and Signac^39^ (v1.14.0). The top and bottom 10% of nFeature_RNA and nFeature_ATAC per patient were removed, as well as percent.mt > 1.5 %, TSS.enrichment < 4 and nucleosome_signal > 2, and down-sampled to 130 nuclei per line. RNA layers followed the scRNA pipeline. Peaks were called using MACS3^40^ and further processed using TF-IDF, FindTopFeatures, SVD and LSI, with standard Signac functions. Modalities were integrated with weighted-nearest-neighbour UMAP (FindMultiModalNeighbors). JASPAR 2024 motifs were added and chromatin deviations calculated with chromVAR 1.26.0.

### CUT&RUN LoV-U experiments

CUT&RUN Low Volume Urea, library preparation and sequencing were performed as described in Zambanini *et al.*^24^ with the following modifications. Organoids were harvested, removed from BME with dispase, dissociated to single cells with TrypLE, washed, and pelleted for 5 minutes at 800 g. Cells were resuspended in 2 ml nuclear extraction (NE) buffer (for 20 ml: 400 µl HEPES-KOH [1 M] pH 8.2, 200 µl KCl [1 M], 5 µl spermidine [2 M], 10 µl IGEPAL 100%, 8 ml glycerol 50%, 20 µl PMSF [1000X], ddH2O to 20 ml), pelleted again, and washed a total of three times. Nuclei were finally resuspended in 40 µl NE buffer per sample. Nuclei pellets were frozen in an isopropyl chamber and stored at -80 °C until processed. CUT&RUN wash buffers were supplemented with 0.025% digitonin and 0.05% BSA. Antibodies used included anti-β-catenin (antibodies-online, ABIN2855042), anti-LEF1 (antibodies-online, ABIN1680678), anti-TCF7L2 (Cell Signalling Technologies, #2569S), and rabbit IgG isotype control (Invitrogen, #100500C). After pAG-MNase digestion, the reaction was stopped with 3 ul of EDTA/EGTA 250 mM mix, and fragments were released by the addition of 2 ul 5M NaCl followed by heating to 37 degrees for 30 min. The supernatant was saved, and the beads resuspended in 50 ul 1X Urea buffer for the second release step. After 30 min incubation at RT, the urea release was combined with the initial supernatant. Samples were subjected to proteinase K digestion and SDS treatment before being purified using phenol chloroform isoamyl alcohol followed by ethanol precipitation. Samples were sequenced to a depth of 5 – 15 million reads on a NextSeq 550 with 36 bp pair-end reads.

### CUT&RUN data analysis

Reads were trimmed with bbmap bbduk^41^ (version 38.18) to remove adapters and poly [AT], [G] and [C] repeat sequences. Reads were aligned to the hg38 genome with bowtie2^42^ (version 2.4.5) with the options –local –very-sensitive-local –no-unal –no-mixed -no-discordant –phred33 –dovetail -I 0 -X 500. Samtools^43^ (version 1.11) suite was used to remove duplicate and incorrectly paired reads. Bedtools^44^ (version 2.30.0) was used to remove reads mapped to the CUT&RUN hg38 suspect list^45^ from bam files. Bigwigs were created for visualization using deepTools^46^ (version 3.5.1-0) with the options -e and -RPCG. After quality control of individual datasets, biological duplicates were merged using samtools. Peaks were called on merged datasets using GoPeaks^47^ with the options -r 8 and the default threshold of adj. p < 0.05, against the corresponding negative control. Wnt peaks were defined for each organoid subtype by combining peaks from β-catenin, LEF1, and TCF7L2 for each organoid line, and then determining regions called in at least 2 of 3 lines. PePr^26^ was used to define differential peaks, inputting the individual biological replicates and negative controls, with the settings p < 0.00005 (default) and subsetting for a calculated FC > 2. Final peak sets were generated after subtracting peaks that consisted of > 50% soft-masked or marked repeat regions. For figure visualization, bigwigs of all three organoid lines per subtype were averaged using deepTools bigwigAverage. Heatmaps and signal intensity plots were created with deepTools computeMatrix and plotHeatmap. Motif analysis was done using HOMER^48^ (version 4.11) findMotifsGenome to find motifs in the hg38 genome using -size 200, and annotatePeaks for motif searching and identification. Peak set gene annotation was done using GREAT^49^ (version 4.0.4) with default parameters. Enrichr^50^ was used for pathway analysis with KEGG and/or Bioplanet pathways, using the entire set of expressed genes according to the scRNA-seq as a background set. Public data for β-catenin in HCT116^27^, and LEF1 and TCF7L2 in HEK293T^24^, K562 (ENCSR343ELW and ENCSR888XZK) and HEK293 (ENCSR240XWM and ENCSR000EUY) were downloaded and compared using the bigwig and peak files.

### Patient tissue samples

This research project complies with all relevant ethical regulations. This project was approved by the Princess Máxima Center Biobank and Data Access Committee (PMCLAB2020-107). Fresh tumor specimens were obtained from surgical resections performed at the University Medical Center Groningen (UMCG) with written informed consent from patients and/or their legal guardians.

### Patient tissue single cell RNA/ATAC sequencing

Single-cell ATAC and gene expression samples and libraries were prepared using the 10x Genomics Chromium Next GEM Single Cell Multiome ATAC + Gene Expression workflow, following the manufacturer’s instructions (CG000375 Rev C and CG000338 Rev F). Cryopreserved tissue fragments were thawed, washed in ice-cold PBS with 5% FBS, and minced to ∼1–2 mm2 if needed. Nuclei were released by 5 min NP-40–based lysis with dounce homogenization on ice, filtered through 70 μm cell strainers (Greiner Bio-One EASYstrainer, Thermo Fisher), and pelleted at 4 °C. Pellets were resuspended in PBS with 2% BSA and Protector RNase inhibitor (Sigma-Aldrich). Intact nuclei were enriched by sorting on a Sony SH800S cell sorter with 100 μm nozzle for 7-AAD (Invitrogen) positive singlets and collected into BSA-coated tubes containing Protector RNase inhibitor. Nuclei were permeabilized, washed, counted on an automated counter (Countess II cell counter, Thermo Fisher) or hemocytometer, and adjusted to ∼600–700 nuclei/μl for loading on a Chromium Controller. Libraries were sequenced on a NovaSeq6000 System (Illumina) with sequencing settings recommended by 10x Genomics and processed using Cell Ranger ARC (v2.0.0).

### Patient tissue single cell RNA/ATAC analysis

We imported 10x Genomics Single Cell Multiome matrices with Read10X_h5, created a Seurat object from RNA counts, and added ATAC data with Signac using hg38 and EnsDb.Hsapiens.v8637–40. ATAC peaks were called with MACS341, peaks overlapping the hg38 unified blacklist were removed, and a peaks assay was built from fragments. SNP-based demultiplexing was performed using Python packages cellsnp lite (v1.2.2) and Vireo (v0.2.3). Genotyping references of the donors were obtained from bulk RNA-sequencing data of the tumor biopsy samples generated in the diagnostic setting. Entries labeled “doublet” or “unassigned” were excluded. For each sample, we trimmed the bottom and top 10 percent of cells by nFeature_RNA and nFeature_ATAC, then applied global filters of mitochondrial RNA fraction below 5 percent, TSS enrichment above 3, and nucleosome signal below 3. ATAC data were processed with TF-IDF, feature selection, and SVD, then corrected across reactions with Harmony36 in LSI space. RNA data were split by reaction and normalized with SCTransform; features correlated with cell cycle or mitochondrial signal were removed using a custom correlator together with S and G2M gene sets, after which PCA was computed. Layers were integrated with IntegrateLayers (method = HarmonyIntegration) to obtain a shared Harmony-corrected PCA space (“harmony.pca”). Embryonal (E) and fetal (F) tumor populations for both patients were defined by analyzing each tumor separately after SCTransform and PCA at low clustering resolution, then assigning states based on canonical markers, with *LEF1* high indicating embryonal and *ALB* and *HNF4A* high indicating fetal. These assignments yielded four groups, 13E, 13F, 17E, and 17F, which were annotated to the full object. We downsampled the object to 300 cells per group, re-ran Harmony on LSI and PCA with 10 dimensions, and visualized *HNF4A*, *LEF1*, and *TCF7L2* on the Harmony PCA. Differential gene expression between embryonal and fetal populations was performed in Seurat using FindAllMarkers on the RNA assay.

### Western blot

Organoids were released from their matrix using dispase and lysed in RIPA Lysis Buffer System (Santa Cruz). After quantification using the BCA protein Assay Kit (Thermo Fisher), protein samples were separated by sodium dodecyl sulfate-polyacrylamide gel electrophoresis (SDS-PAGE). Proteins were then transferred onto polyvinylidene difluoride (PVDF) membranes (Millipore) using semi-dry method (Trans-blot Turbo transfer system). Membranes were blocked in 5% BSA (Sigma-Aldrich) for 1 h at room temperature and incubated with anti-TCF7L2 (1:1,000, Cell Signaling Technologies, #2569), anti-LEF1 (1:1,000, Cell Signaling Techology, #2230), and GAPDH (1:2,000, Cell Signaling Technology, #2118) antibodies, followed by incubation with IRDye 800CW goat anti-rabbit IgG secondary antibody (1:5,000, LI-COR) and IRDye 680RD goat anti-mouse IgG secondary antibody (1:10,000, LI-COR). Immunoreactive proteins were subsequently visualized using the Bio-rad ChemiDoc Imaging System.

## Supplementary figures

**Supp. Figure 1.**
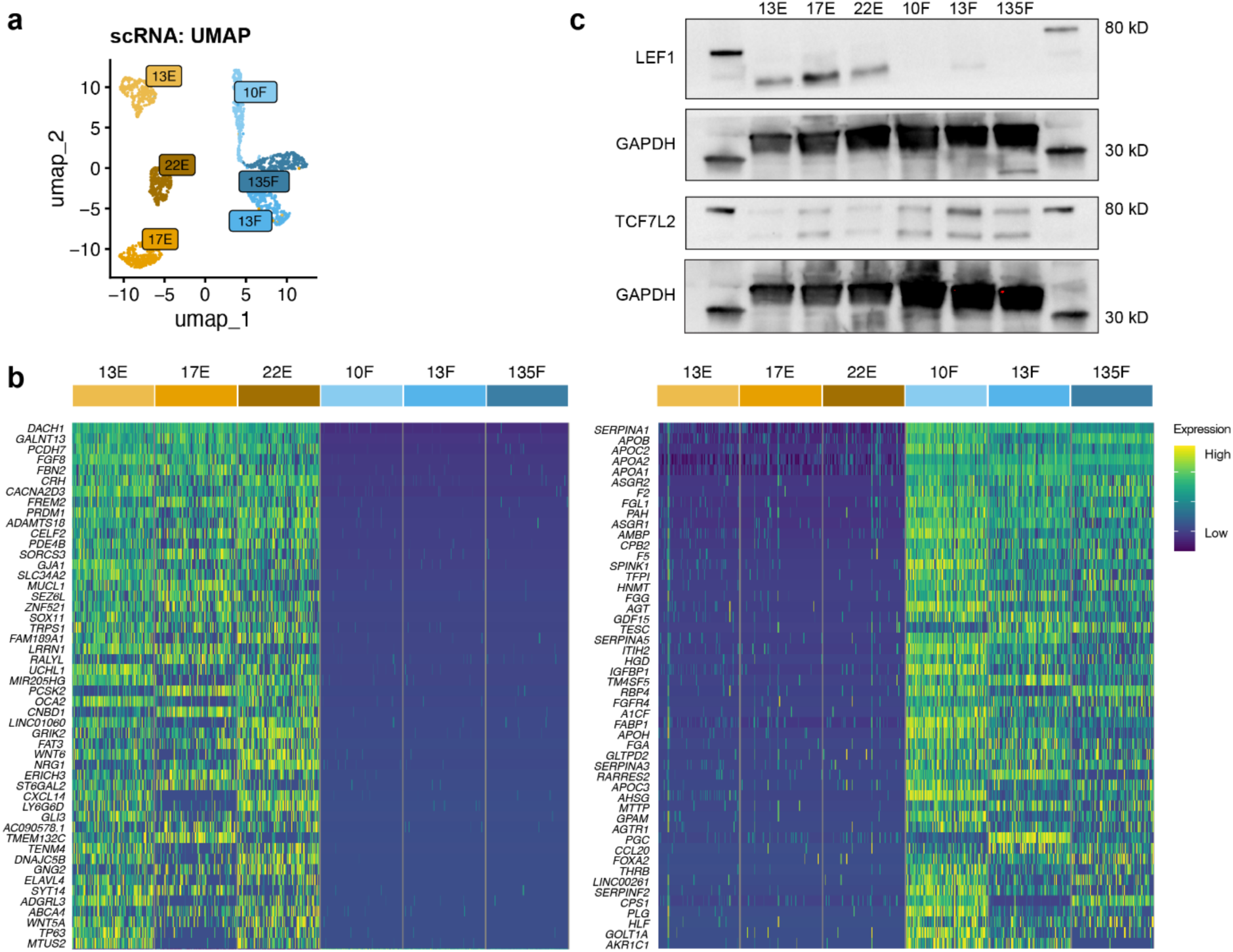
Organoid scRNA analyses. a. scRNA-seq UMAP of E and F tumor organoid models. b. Heatmap showing the top 50 differentially expressed genes in E and F organoids. c. Western blots showing LEF1 and TCF7L2 in all organoid models with their respective GAPDH loading control.

**Supp. Figure 2.**
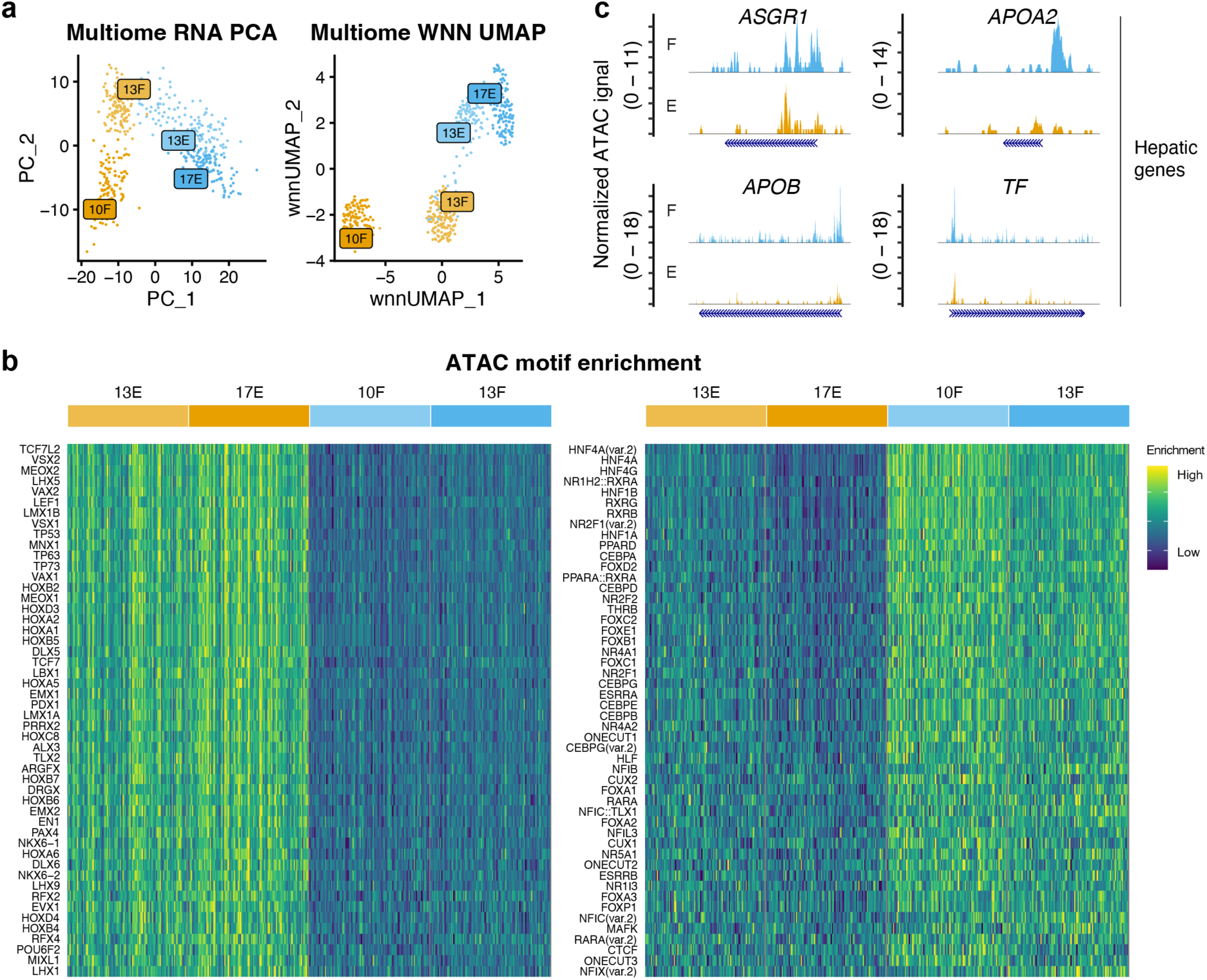
Chromatin accessibility analysis of PDOs. A. scRNA/ATAC-seq PCA and Weighted Nearest Neighbor (WNN) UMAP of E and F tumor organoids. B. Heatmap showing differentially enriched motifs in E and F organoids. C. Coverage plots showing ATAC signal at promoter regions of selected hepatic targets split for E and F organoids.

**Supp. Figure 3.**
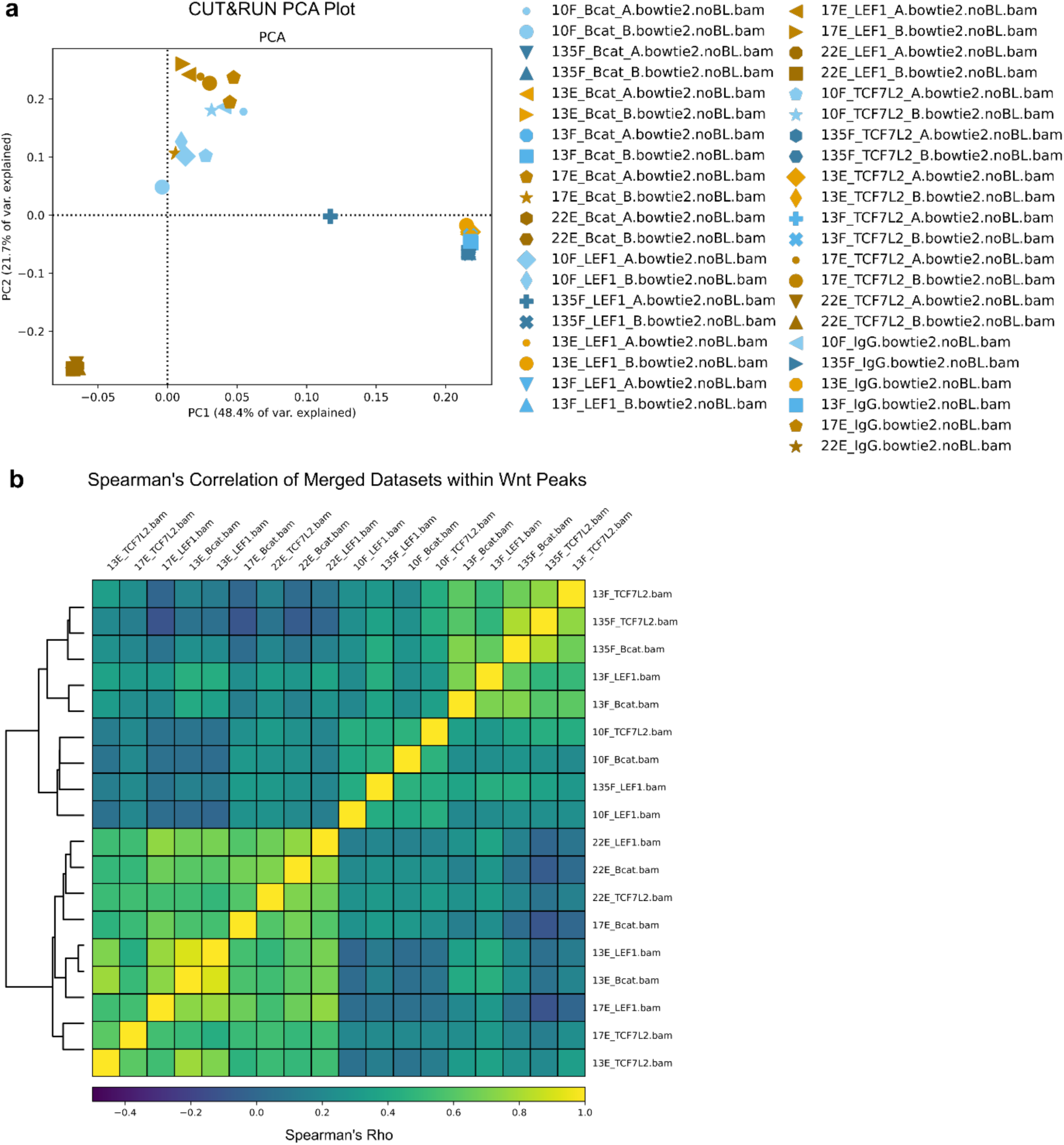
CUT&RUN LoV-U of PDOs. A. PCA plot after genome-wide binning of individual replicates, showing mainly separation based on genotype, due to karyotypic differences between the organoid lines. B. Spearman’s correlation heatmap showing correlation of merged replicates within Wnt peaks. Two main unsupervised hierarchical clusters are formed, separating E and F lines.

**Supp. Figure 4.**
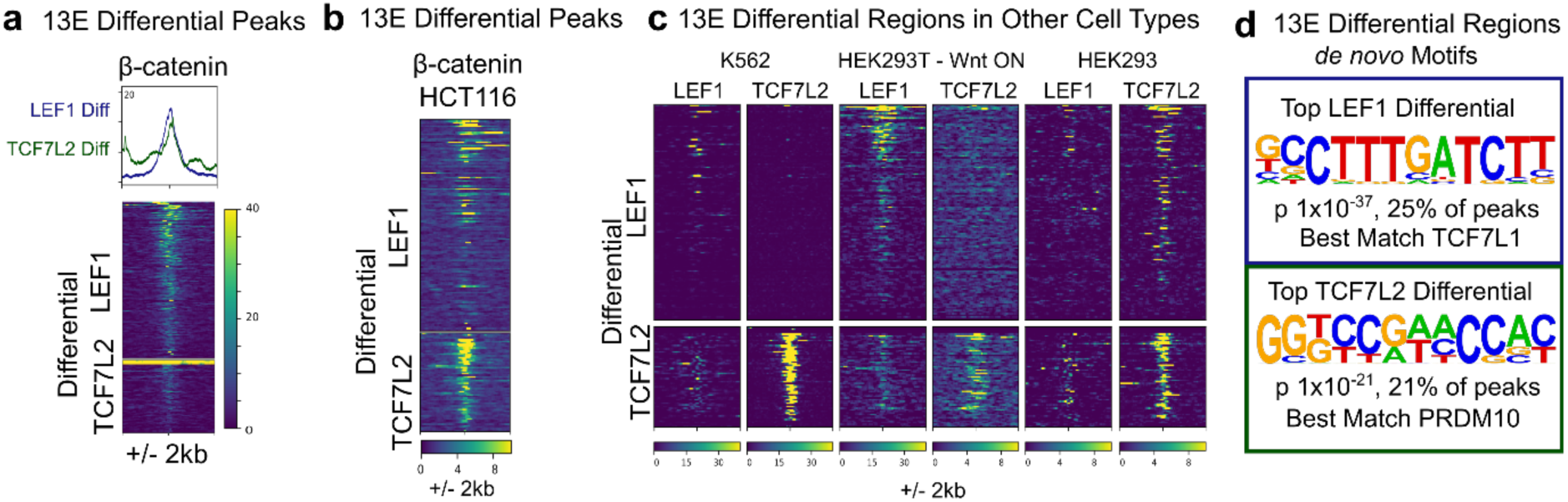
Differential LEF1 and TCF7L2 occupied loci in 13E. a. Heatmap showing β-catenin CUT&RUN signal within 13E differential regions, showing comparable average signal strength. b. Heatmap showing β-catenin CUT&RUN signal on 13E differential regions in *TCF7L2*-expressing HCT116 cells, showing higher occupancy in the TCF7L2 peaks. c. Heatmaps of LEF1 and TCF7L2 signal in published matched ChIP and CUT&RUN datasets of cells expressing both LEF1 and TCF7L2, on the 13E differential regions. d. Top *de novo* motifs identified in the LEF1 and TCF7L2 differential regions.

**Supp. Figure 5.**
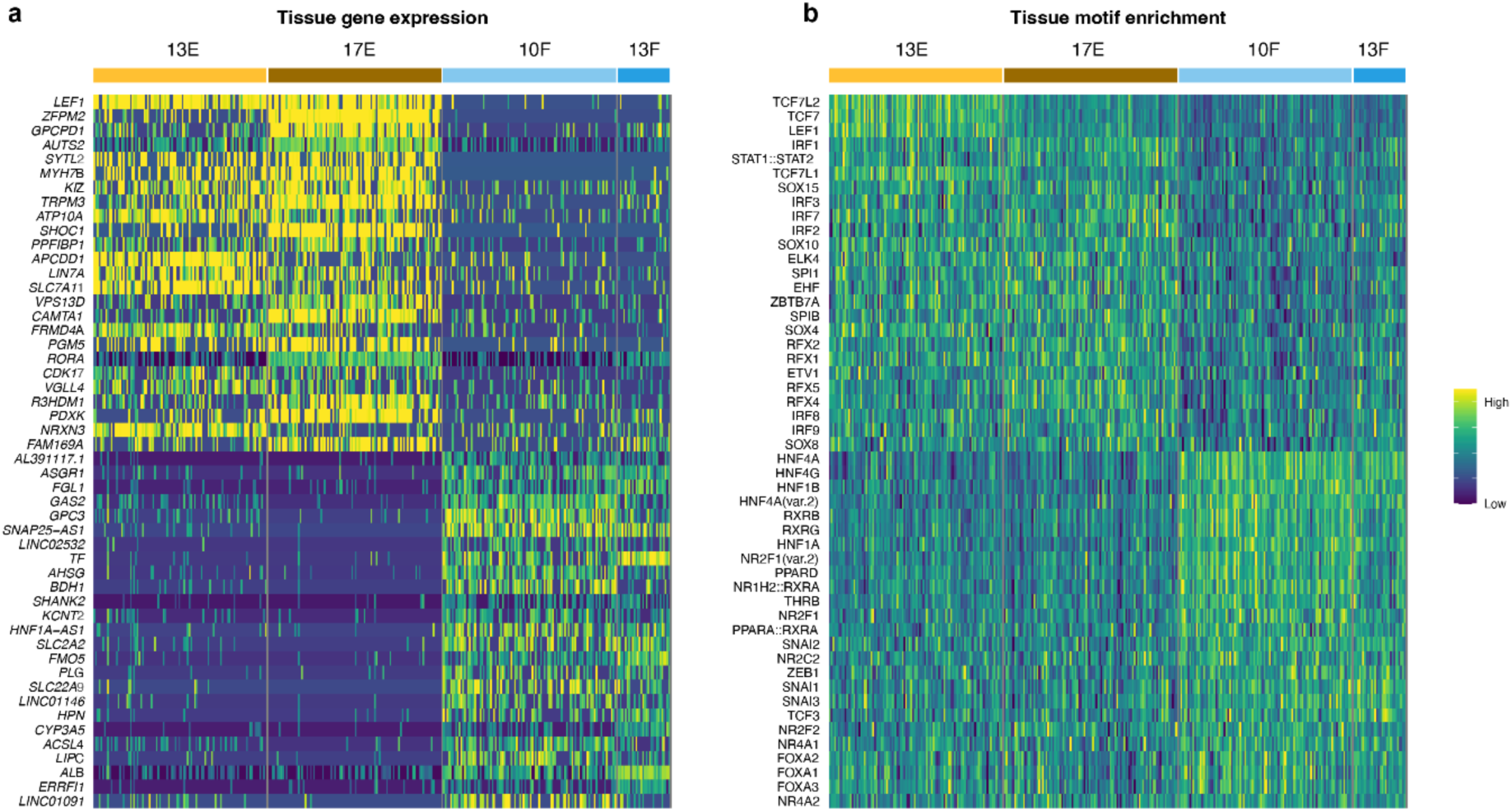
TCF7L2 directs a hepatic lineage gene program. a. Heatmap showing the top differentially expressed markers between the E and F subtypes in patient tissue samples from patients 13 and 17. b. Heatmap showing the top differentially enriched motifs in ATAC peaks between the E and F subtypes in patient tissue samples from patients 13 and 17.

